# Efficient growth factor production stabilizes cooperation in a cellular public goods game

**DOI:** 10.64898/2026.10.29.762779

**Authors:** Jin Eun Shin, Justin Pritchard, Marco Archetti

## Abstract

Eukaryotic cells secrete growth factors that facilitate the proliferation and survival of neighboring cells in a coordinated manner essential for tissue development, repair, and homeostasis. However, mutant cells that fail to produce these factors may exploit the benefits provided by neighboring producers, thereby gaining a proliferative advantage that could lead to the extinction of the producer clone. Evolutionary theory predicts that preventing this “tragedy of the commons” requires a fixed spatial structure or other mechanisms that enable cells to interact preferentially with copies of themselves. Contrary to this expectation, we found that, in competition experiments, cells producing the diffusible growth factor interleukin-3 (IL3) resist invasion by non-producing mutant cells even in well-mixed populations, and that non-producers are driven to extinction instead. Using evolutionary game theory, we show that this outcome arises when the cost-to-benefit ratio of growth factor production falls below a critical threshold. We derive this tipping point analytically across the full parameter space of benefit function shape and group size and confirm that IL3 production in our experimental system satisfies this condition — with a production cost less than 0.1% of the benefit conferred. These results demonstrate that efficient public good production alone can resolve the tragedy of the commons in populations of cells, and potentially in other biological systems.

## Introduction

Eukaryotic cells communicate via growth factors that can direct proliferation during development and that regulate growth, homeostasis and repair in adult tissues. (Massagué, 1998; Gurdon & Bourillot, 2001). Growth factors are commonly secreted and act in a diffusible manner, binding to specific cell surface receptors to trigger downstream signaling cascades that can promote cell growth and survival (Heldin, 1995; Lemmon & Schlessinger, 2010). As a result, most growth factors exert paracrine effects — stimulating not only the cells that produce them but also neighboring cells (Böttcher & Niehrs, 2005). In the terminology of evolutionary theory, this constitutes cooperative behavior at the cellular level (West, Griffin, & Gardner, 2007; Crespi, 2001). Although autocrine and nondiffusible signaling modes also exist (Schlessinger, 2000), paracrine signaling is of particular importance because it coordinates growth across cells — a function that is essential during tissue development and homeostasis (Lander, 2011).

If growth factors acted only through autocrine signaling, mutations that increase production could spread clonally and drive unbalanced, localized overgrowth. Paracrine signaling constrains this dynamic: because the benefit of excess growth factor is shared with neighboring cells that bear none of the production cost, overproducing mutants cannot translate their investment into a private fitness advantage (Pepper, 2012; Archetti, Ferraro, & Christofori, 2015). The benefit is, in effect, diluted across the population. Paracrine growth factor sharing therefore promotes uniform growth and suppresses the invasion of overproducing mutants — functioning as an evolutionary brake on runaway clonal expansion (Frank, 2010).

This, however, raises the opposite problem: mutant cells that produce little or no growth factor could spread by free-riding on the growth factors supplied by surrounding producers. As non-producers increase in frequency, the shared growth factor becomes depleted, ultimately collapsing the population’s cooperative capacity — a “tragedy of the commons” (Hardin, 1968) at the cellular level (Axelrod, Axelrod, & Pienta, 2006, Archetti & Pienta 2019). How do cells avoid this outcome? Evolutionary theory has traditionally explained the maintenance of costly cooperative behaviors, such as public good production, by invoking positive assortment (Hamilton, 1964): when individuals interact preferentially with others of the same type, non-producers are disproportionately surrounded by other non-producers and thus receive less benefit from the public good. In cell populations, such assortment can arise from kin recognition or, more commonly, from the fixed spatial structure of solid tissues, where cell interactions are inherently local (West, Griffin, & Gardner, 2007; Fletcher & Doebeli, 2009; Kerr et al., 2002; Nadell, Foster, & Xavier, 2010).

Here we report that producer cells growing in suspension — that is, in a well-mixed population with no spatial structure or kin assortment — not only resist invasion by non-producers but consistently drive them to extinction, regardless of initial frequency. This outcome is apparently surprising given the logic of the “tragedy of the commons”, which predicts that producers should be vulnerable to free-riders in the absence of positive assortment. To understand this result, we combine experimental evolution using genetically modified cell lines with mathematical modelling in the framework of evolutionary game theory. We show that extinction of non-producers can occur without assortment when the cost-to-benefit ratio of public good production falls below a critical threshold — and that IL3 production in our system satisfies this condition.

## Results

### In a well-mixed population of IL3 producers and non-producers, non-producers go extinct

Ba/F3 is a murine pro-B lymphocyte cell line that requires interleukin-3 (IL3) for survival and proliferation but does not produce it endogenously. Ba/F3 grows in suspension rather than in adherent monolayers, making it a natural model system for well-mixed population dynamics. To generate a producer clone, we engineered Ba/F3 cells to express IL3 via a tagged lentiviral cassette, followed by selection of stably transduced cells. As expected, wild-type non-producer cells did not survive in the absence of exogenous IL3 but proliferated normally when IL3 was supplied exogenously in the growth medium; producer cells remained viable without exogenous IL3, which improved their growth rate only marginally (**Figure 1a**). The negligible benefit of exogenous IL3 supplementation to producer cells indicates that endogenous IL3 production is highly efficient — nearly saturating the available proliferative benefit.

**Figure 1.**
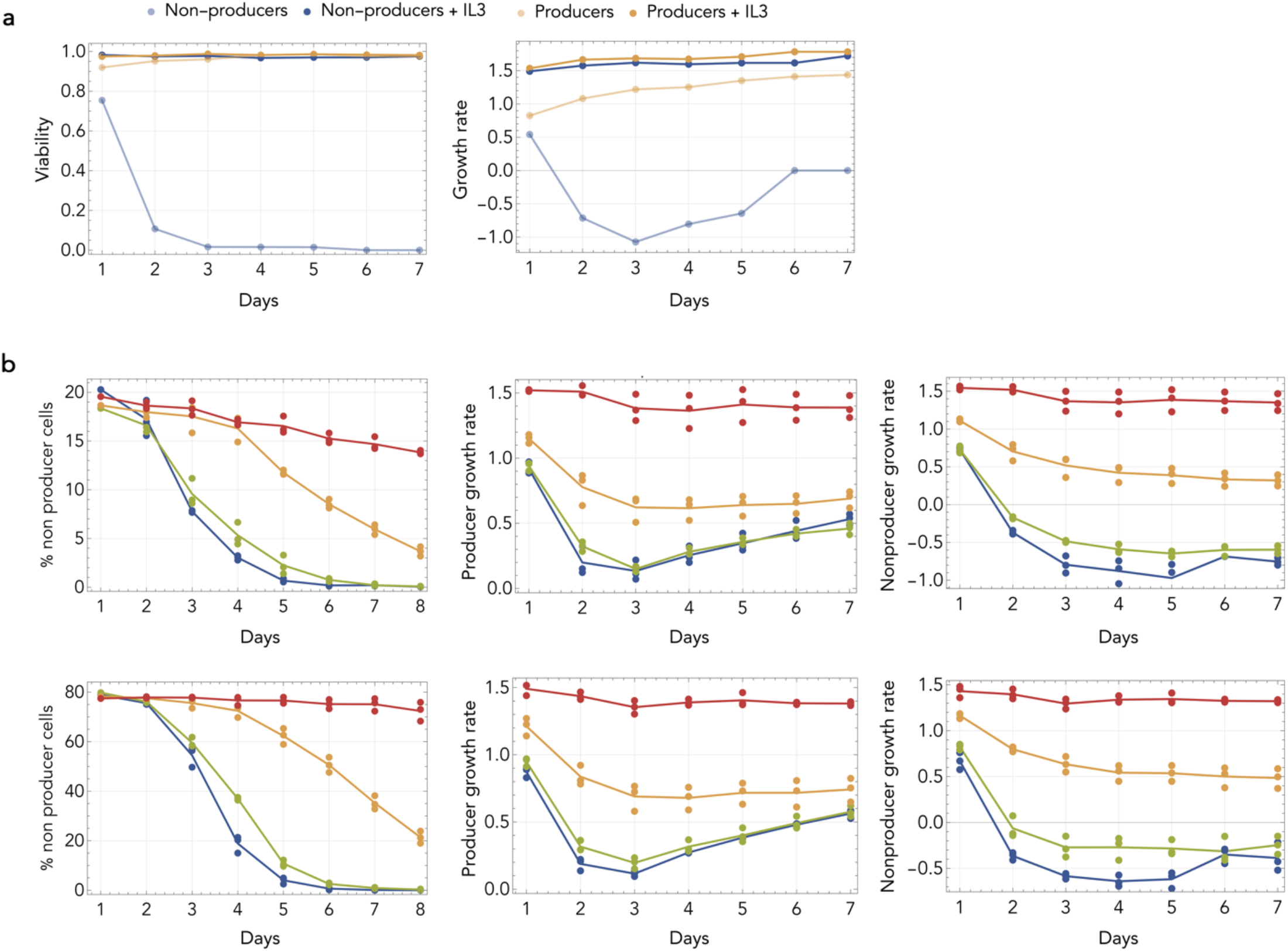
Competitive dynamics of IL3 producer and non-producer Ba/F3 cells. **a**: Growth rates and viability of pure producer and non-producer populations cultured with or without exogenous IL3 over 7 days. **b**: Mean frequency and growth rate dynamics in co-cultures initiated at 20% (top) or 80% (bottom) non-producers, measured over 8 days across different IL3 conditions (no IL3, blue; 0.01 ng ml^−1^ IL3, green; 0.05 ng ml^−1^ IL3, yellow; exogenous IL3 matching producer output, red). Points represent biological replicates.

When producer and non-producer cells were cultured together, non-producer frequency declined consistently over time regardless of initial composition (**Figure 1b**). Supplementing the medium with exogenous IL3 slowed this decline but did not reverse it, and non-producers ultimately went extinct under all conditions tested. The rate of decline was frequency-dependent, with non-producers decreasing more rapidly at lower initial frequencies.

These results show that, in a well-mixed population, producing IL3 does not render the producer clone susceptible to invasion. There is no tragedy of the commons in this system. Instead, it is the non-producer clone that is driven to extinction. To understand how the logic of the “tragedy of the commons” can be reversed in the absence of positive assortment, we examined the problem using evolutionary game theory.

### Evolutionary game theory identifies two routes to the extinction of non-producers

We model the population as a public goods game in which producers and non-producers interact in groups of size *N*, and the benefit of IL3 is a function of the number of producers in the group. We adopt the standard framework used in evolutionary game theory to study cooperation for the production of public goods by multiple players, with an important extension: we allow the benefit of the benefit of the growth factor to be a nonlinear function of the number of producer cells.

When the benefit function is nonlinear — as it is in our system — the equilibria of the replicator dynamics cannot be located analytically. However, by leveraging properties of Bernstein polynomials, the dynamics can be precisely characterized and reliable approximations for the equilibria derived by analysing the Bernstein coefficient of the gradient of selection (Archetti 2018). Depending on the cost of production *c* relative to the benefit function, five qualitatively distinct outcomes are possible (**Figure 2b–c**). If *c* ≥*β*_MAX_ there is a unique stable equilibrium *x* = 0 (producers go extinct). If Max[Δ*b*_0_, Δ*b*_*N*_] ≤ *c* < *β*_MAX_ there are two stable equilibria: a pure equilibrium *x* = 0 made of all non-producers and a mixed equilibrium *x*\*; the basins of attraction of these two equilibria are separated by a mixed unstable equilibrium *x*^; if the initial frequency of producers *x* < *x*^ the population will evolve to *x* = 0; if *x* > *x*^ it will evolve to *x*\*. If Δ*b*_0_ ≤ *c* < Δ*b*_*N*_ (this case can exist only for *h* > 0.5) there is a unique interior unstable equilibrium *x*^ separating the basins of attraction of two pure stable equilibria: *x* = 0 and *x* = 1; if *x* < *x*^ the population will evolve to *x* = 0; if *x* > *x*^ it will evolve to *x* = 1. If Δ*b*_*N*_ ≤ *c* < Δ*b*_0_ (this case can exist only for *h* < 0.5) there is a unique interior stable equilibrium *x*\*, to which the population will converge irrespective of the initial frequencies of the two types. If *c* < Min[Δ*b*_0_, Δ*b*_*N*_] there is a unique stable equilibrium *x* = 1 (non-producers will go extinct).

**Figure 2.**
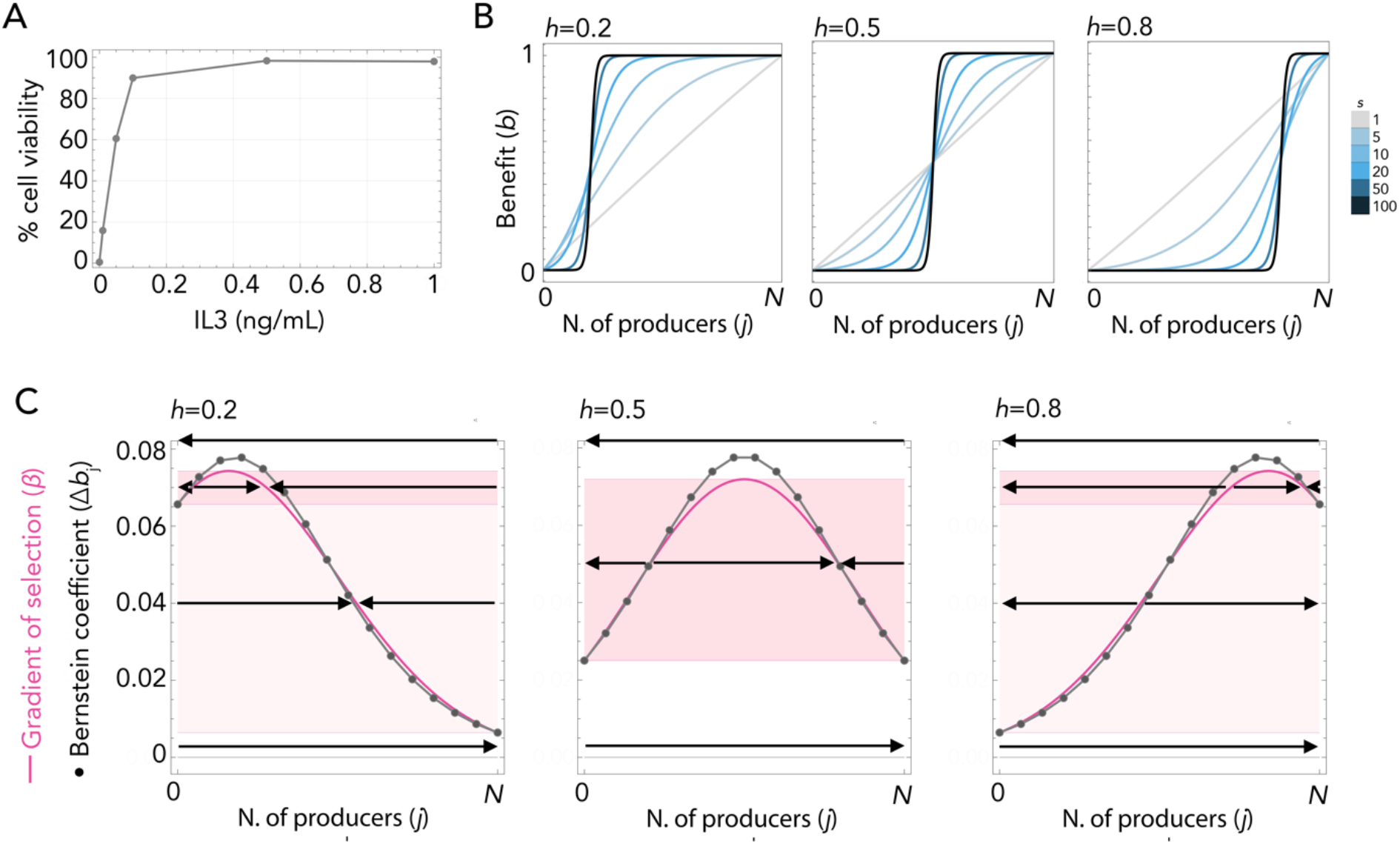
Nonlinear IL3 benefit functions and their evolutionary dynamics. **A:** Cell viability as a function of exogenous IL3 concentration. **B:** Benefit functions as a function of the number of producers in a group of size *N*, shown for three values of the inflection point *h* (columns: *h* = 0.2, 0.5, 0.8) and seven values of the steepness parameter *s* (line shading, light to dark). **C:** Gradient of selection (solid line) and Bernstein coefficients (points) of the replicator dynamics for each value of *h* in **B**, assuming *N* = 20 and *s* = 5. Arrows indicate the direction of evolutionary change. Shaded regions indicate cost-to-benefit ratio ranges corresponding to qualitatively distinct dynamical regimes (dark blue: two interior equilibria; light blue: one interior equilibrium).

These results reveal two distinct routes by which non-producers can be driven to extinction. The first — which we term the “*low-cost hypothesis*” — occurs when the cost of production *c* is sufficiently small relative to the benefit, placing the system in the last case listed above regardless of benefit shape. The second — the “*convex-benefit hypothesis*” — occurs when the inflection point *h* of the benefit function is high (*h* > 0.5), meaning that the benefit of IL3 accelerates with increasing producer frequency (increasing returns); this places the system in the third case in our list above, where non-producers go extinct from high initial producer frequencies. To determine which of these mechanisms operates in our system, we next estimate the shape of the IL3 benefit function and the cost of its production.

### Parameter estimation reveals diminishing returns and negligible production cost

The cost of IL3 production can in principle be estimated by measuring the growth rates of producers and non-producers in the presence of saturating exogenous IL3: under these conditions, both cell types receive the same benefit, thus any remaining fitness difference reflects the cost borne by producers alone. In our system, however, no detectable growth rate difference was found between the two clones under saturating IL3 (**Figure 1a**). This is consistent with the low-cost hypothesis but does not by itself distinguish between the two routes to non-producer extinction, since a concave benefit function could also produce this pattern.

To resolve this ambiguity, we estimated the shape of the benefit function directly by fitting observed population growth rates to the predicted average fitness of mixed populations. Critically, note that it is not sufficient to estimate the benefit function from the dose–response curve of IL3 concentration alone, because the same sigmoidal concentration– response curve is consistent with the benefit function being in either its concave or convex regime depending on where the operating range of producer frequencies falls. We measured the growth of mixed producer–non-producer populations two days after seeding at a range of initial ratios and fit these data to the model’s average fitness function across a range of group sizes *N* (**Figure 3a**). Although *N* — the number of cells within the diffusion range of IL3 — cannot be measured directly, the estimated parameters are robust across a wide range of *N* values and so do not depend critically on this unknown (**Figure 3a–b**).

**Figure 3.**
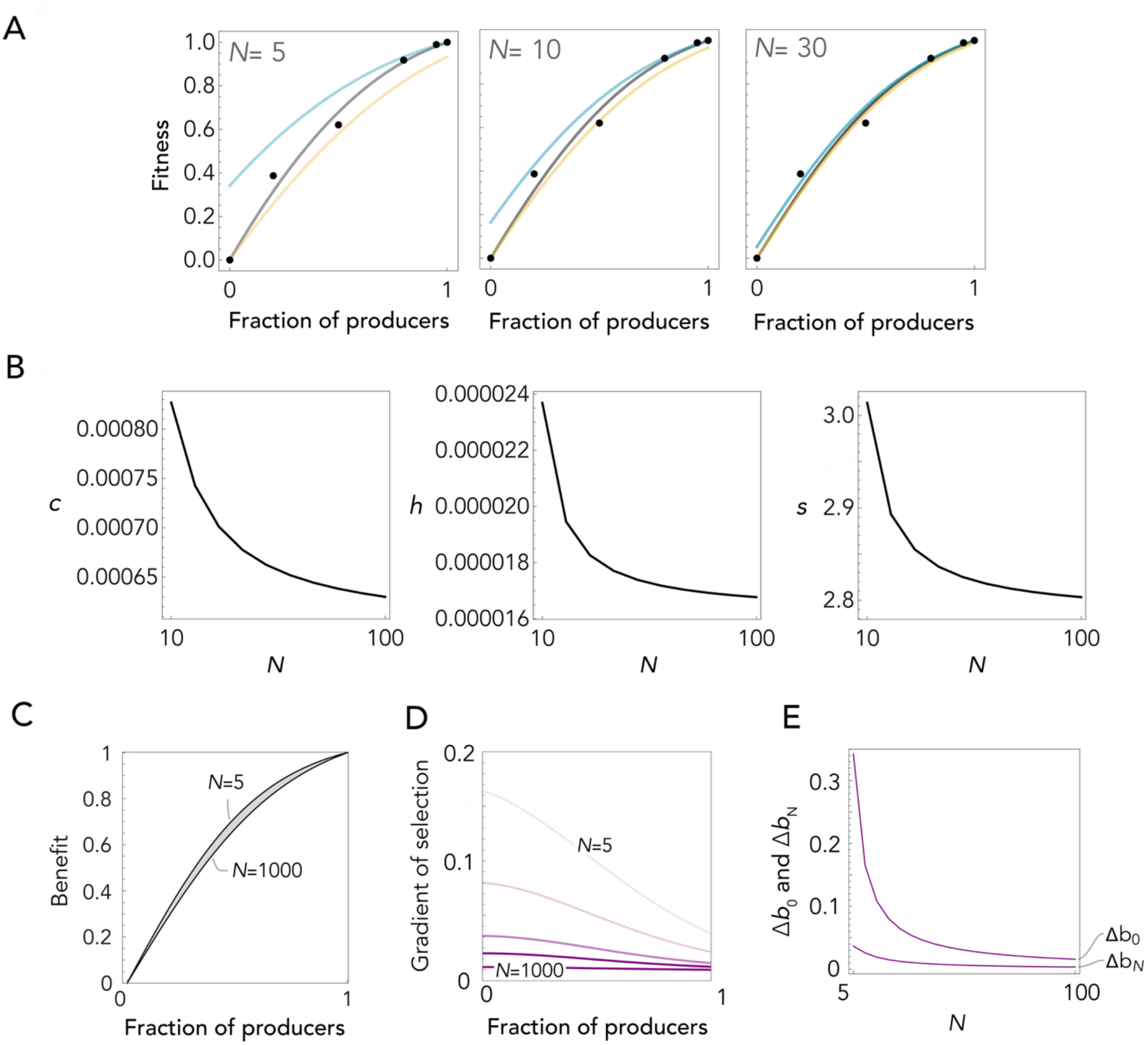
Parameter estimation for the IL3 public goods game and the tipping point for non-producer extinction. **A:** Observed growth rates of mixed producer–non-producer populations (black points, mean of two measurements) and model-predicted average fitness of the mixed population (black curve), producers (blue curve), and non-producers (yellow curve), as a function of producer fraction, for *N* = 5, 10, and 30. **B:** Estimated model parameters *c, s*, and *h* (assuming maximum benefit *b* = 1) as a function of group size *N*, obtained by fitting model-predicted average population fitness to observed growth rates in **A. C:** Benefit functions reconstructed from the estimated parameters in **B** for *N* ranging from 5 to 1000 (light to dark). **D:** Gradient of selection as a function of producer fraction for *N* ranging from 5 (light) to 1000 (dark). **E:** Δ*b*_0_ and Δ*b*_N_ as a function of *N* for the estimated parameters. The tipping point — the maximum cost-to-benefit ratio *c* below which non-producers go extinct — is given by Min[Δ*b*_0_, Δ*b*_N_].

The fitted parameters reveal that the benefit of IL3 is a concave function of producer frequency (*h* ≈ 0, indicating strongly diminishing returns) with a shallow slope (*s* ≈ 3; **Figure 3b–c**). The estimated production cost is *c* < 0.001 relative to a maximum benefit normalized to 1 — less than 0.1% of the total benefit conferred by IL3 (**Figure 3b**). Together, these parameter estimates place our system firmly in the low-cost regime: the cost of IL3 production is far below the tipping point threshold, not because of convex benefits, but because the cost-to-benefit ratio is extremely low – production of the public good is exceptionally efficient.

Given these estimates, the replicator dynamics predicted by the model drives non-producers to extinction for group sizes up to at least *N* = several hundred (**Figure 3d**). The tipping point — the maximum cost-to-benefit ratio below which non-producers go extinct — is approximately 0.0035 for *N* = 100 and falls below 0.001 for *N* > 500 (**Figure 3e)**. Since the estimated cost lies well below this threshold across all biologically plausible values of *N*, the model consistently predicts extinction of non-producers in our system.

### The tipping point for non-producer extinction across the full parameter space

The low-cost condition derived above applies not only to our IL3 system but defines a general criterion for the extinction of non-producers in any nonlinear public goods game. To map this criterion across the full parameter space, we calculated the tipping point Min[Δ*b*_0_, Δ*b*_*N*_] — the maximum cost-to-benefit ratio below which non-producers go extinct — for all combinations of benefit shape (*h, s*) and group size *N* (**Figure 4**). Three patterns emerge consistently. First, the tipping point decreases with increasing group size *N*: larger groups dilute the per-capita benefit of each producer, making it harder for low-cost production alone to confer a net advantage. Second, the tipping point decreases as the steepness *s* of the benefit function increases: shallower benefit functions maintain a producer advantage across a wider range of producer frequencies, enlarging the parameter regime in which non-producers go extinct. Third, the tipping point is highest for sigmoid benefit functions (intermediate *h*) and lower for both strongly concave (*h*→0) and strongly convex (*h*→1) functions.

**Figure 4.**
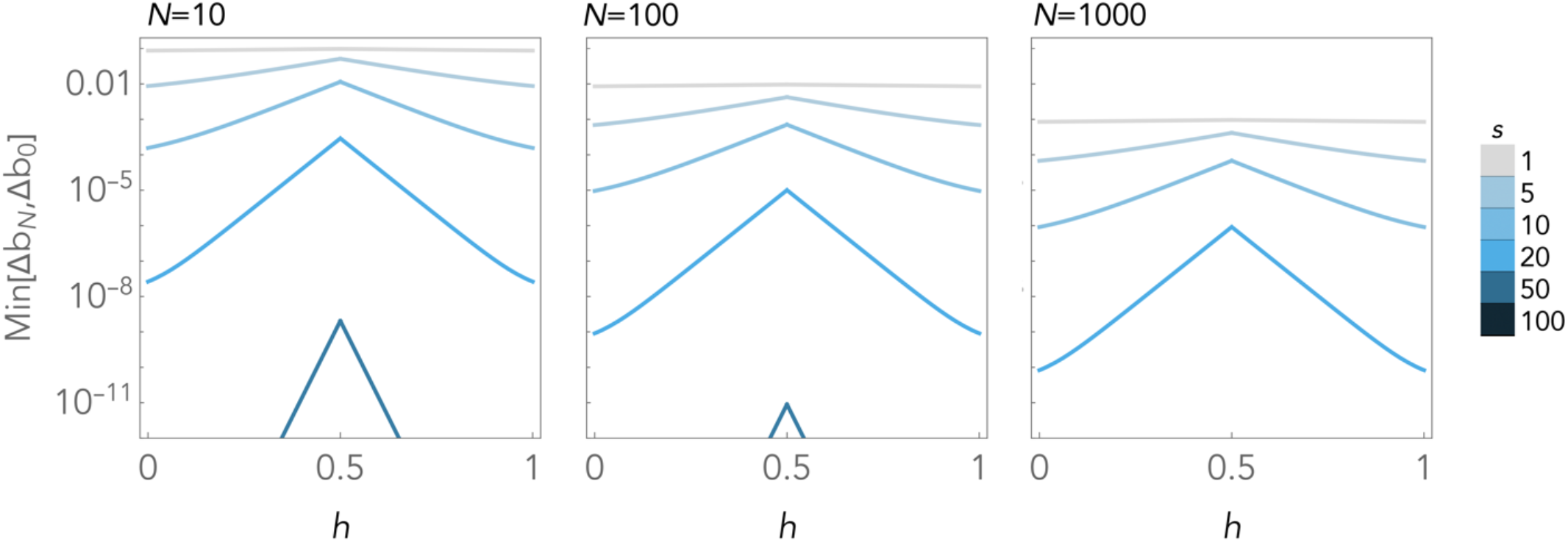
Tipping points for non-producer extinction across the full parameter space. The tipping point Min[Δ*b*_0_, Δ*b*_*N*_] — the maximum cost-to-benefit ratio below which non-producers go extinct — is shown as a function of the benefit function inflection point *h*, for six values of the steepness parameter *s* (line shading, light to dark) and three group sizes *N* = 10, 100, and 1000 (panels left to right).

## Discussion

We have shown that in a well-mixed population — lacking spatial structure, preferential interaction between kin or other forms of positive assortment — an engineered IL3-producing Ba/F3 clone consistently drives a non-producing clone to extinction, regardless of initial frequency. This outcome contradicts the common belief about public goods that producers should either go extinct through a tragedy of the commons or coexist with non-producers at a mixed equilibrium. Neither occurs in our system. Instead, the very act of producing a diffusible growth factor confers a net competitive advantage on producers.

We identified two conditions under which non-producers can be driven to extinction without positive assortment: when production cost is sufficiently low (the low-cost hypothesis), or when the benefit function is convex — that is, when producers exhibit increasing returns (the convex-benefit hypothesis). By fitting the model to experimental growth rate data, we established that, in our case, the IL3 benefit function is in fact concave (strongly diminishing returns, *h* ≈ 0), ruling out the convex-benefit hypothesis. The production cost, meanwhile, is less than 0.1% of the maximum benefit — well below the tipping point threshold that permits non-producer extinction. The proximate cause of extinction in our system is therefore low cost, not benefit convexity. Finally, we derived this tipping point analytically across the full parameter space of benefit shape and group size, showing that it decreases with increasing *N*, with increasing steepness *s*, and when benefits are strongly concave or convex rather than sigmoid.

### The theoretical possibility of cost-driven extinction has not previously been demonstrated empirically

The canonical framework for studying public good production is the N-person Prisoner’s Dilemma (NPD), in which producers pay a cost *c* and all contributions are pooled, multiplied by a reward factor *r* > 1, and redistributed equally to all group members regardless of whether they contributed (Archetti & Scheuring 2012). In this linear model, producers go to fixation only if *r* > *N*; that is, only if the collective reward exceeds the group size. Because this condition is generally considered unrealistic for large groups, the standard conclusion has been that cooperation in public goods games requires positive assortment to be maintained (Fletcher & Doebeli, 2009; West, Griffin, & Gardner, 2007). The NPD, however, rests on the assumption that each producer’s contribution has an additive, linear effect on the public good — an assumption that is biologically unrealistic for secreted molecules. Growth factors, like most signaling ligands and most biological molecules, exhibit nonlinear dose–response relationships: their effects saturate at high concentrations and may accelerate or decelerate depending on receptor occupancy and downstream signaling dynamics (Lemmon & Schlessinger, 2010; Böttcher & Niehrs, 2005). IL3 is no exception, as shown by the concave benefit function we estimate for our system.

When the linearity assumption is removed, the dynamics of public goods games change fundamentally. The existence of tipping points in nonlinear public goods games has been noted theoretically (Archetti & Scheuring, 2011), but the possibility of a low enough cost-to-benefit ratio has received little attention as a mechanism for the maintenance of cooperation. Our results provide the first experimental evidence that a public goods game can produce the extinction of non-producers without positive assortment, without frequency dependence, and without convex benefits — simply because the cost of production is negligible relative to the benefit conferred. This finding rehabilitates a theoretically valid but empirically neglected regime of public goods dynamics.

### Why is IL3 production so efficient in our system?

The proximate cause of non-producer extinction in our system is unambiguous: the IL3 benefit function is concave with a very low inflection point (*h* ≈ 0) and shallow slope (*s* ≈ 3), producing slowly diminishing returns that keep the tipping point well above the actual cost of production across all biologically plausible group sizes. What drives this exceptionally low production cost in the first place, however, is not immediately clear.

Two non-mutually exclusive explanations are worth considering. The first is that optimal culture conditions reduce the metabolic burden of transgene expression. Malakar & Venkatesh showed in its study with *E. coli* that burden induced by gene expression on their growth is lower when cultured in resource-rich medium than in resource-deficient medium (Malakar & Venkatesh, 2013). In a similar fashion, in vitro mammalian cell culture provides daily nutrient replenishment and constant environmental conditions, potentially allowing cells to sustain IL3 secretion without the energetic trade-offs that would apply in a resource-limited in vivo environment. If this explanation is correct, the low cost we observe may be partly an artifact of the experimental system rather than a general property of IL3 production.

The second explanation concerns the engineering strategy used to generate the producer clone. When a lentiviral vector is used to introduce a transgene, the resulting cell pool is heterogeneous: individual cells receive different numbers of viral integrations and therefore express the transgene — and bear its metabolic cost — at different levels. As this pool is expanded in culture, clones with lower expression costs will tend to proliferate faster and increase in frequency, so that the producer line used in experiments is enriched for low-cost producers by clonal selection. This process is absent in systems where the non-producer is the engineered line — that is, where non-producers are generated by knocking-out growth factor genes in an established producer background (Archetti, Ferraro, & Christofori, 2015). In those systems, the cost of production reflects the evolutionary history of the producer lineage rather than selection during an engineering protocol, and the higher resulting costs lead to coexistence of producers and non-producers, or even to producer decline over time. The contrast between these outcomes and ours underscores the importance of production cost as the key parameter governing competitive dynamics in growth factor public goods games.

These two explanations are not mutually exclusive and may act together. Distinguishing between them would require comparing production costs across culture conditions and engineering strategies — experiments that would also shed light on how generalizable the low-cost regime is likely to be in natural cell populations.

### How general is the low-cost regime likely to be?

Our results raise a natural question: how often does growth factor production in natural cell populations satisfy the low-cost condition, and is the tragedy of the commons therefore less common at the cellular level than theory has assumed? Two lines of evidence suggest that clonal evolution may lead to low production cost. First, some cells appear to produce paracrine growth factors at levels just sufficient to sustain autocrine signaling and survival, without overactivating downstream pathways (Böttcher & Niehrs, 2005; Archetti & Pienta, 2019; Villarino, 2007) — a pattern consistent with production being tuned to near-minimal cost. Second, the metabolic cost of secreting a signaling protein need not be large under some circumstances. As ligand affinity affects the ligand concentration required to achieve receptor occupancy and signaling, a high-affinity ligand may achieve a level of signaling output with lower ligand concentrations and, in some signaling systems, lower receptor abundance than a low-affinity ligand, potentially reducing the associated synthetic and energetic cost. (Lemmon & Schlessinger, 2010; Heldin, 1995; Wagner, 2005; Hui et al., 2015; Macdonald-Obermann & Pike, 2024; Freed et al, 2017). Whether IL3 or other growth factors fall into this low-cost category in vivo, however, remains to be tested directly.

Several approaches could make this question tractable. The most direct would be to measure the shape of the benefit function — specifically the parameters *h* and *s* — in additional producer–non-producer systems, since the tipping point depends critically on these values. A complementary approach would be to estimate production costs from first principles: knowing the likely secretion rates, ligand concentrations, and receptor expression levels, one could estimate the ATP cost of growth factor production as a fraction of a cell’s total energy budget and compare this to the tipping point derived from the benefit function. At a population level, dependency data from large cancer cell line screens (Tsherniak et al., 2017) could provide a rough estimate of the fitness consequences of growth factor production or receptor signaling loss across diverse cellular contexts, allowing inferred costs to be compared with the thresholds derived from our model.

It is important to acknowledge that the dynamics we describe here operate under idealized conditions — a well-mixed suspension culture with two clonal populations and a single growth factor. Natural cell populations, whether in development or in tumors, are more complex. Spatial structure, stromal interactions, intra-tumor heterogeneity, and the diffusibility of specific growth factors all modulate the effective interaction range and group composition in ways that our model does not capture (Axelrod, Axelrod, & Pienta, 2006; Nadell, Foster, & Xavier, 2010; Archetti & Pienta, 2019; Greaves & Maley, 2012; Marusyk, Almendro, & Polyak, 2012; Pepper, 2012). In solid tissues, spatial structure may reinforce producer advantage through assortment, potentially making the low-cost condition easier to satisfy in vivo than in our well-mixed system. In tumors, by contrast, microenvironmental heterogeneity and selection for rapid growth may elevate production costs or alter benefit functions in ways that push systems away from the extinction regime. Whether the low-cost condition is met in any of these contexts is an empirical question that our theoretical framework is not equipped to address and will require extending our model and experimental system.

Our system is deliberately simplified: IL3 production is driven by an engineered transgene, the population is well-mixed, and only two clonal types compete. These simplifications isolate the mechanism of interest but mean that extrapolation to natural tissues or tumors requires caution and further empirical work. Our results, however, demonstrate that the tragedy of the commons is not an inevitable outcome of cooperation for secreted factors in cell populations. When production costs are sufficiently low, producers can outcompete non-producers even without spatial structure or kin assortment. The Ba/F3–IL3 system provides the first experimental demonstration of this regime, and the tipping point analysis we derive offers a quantitative tool for assessing whether other growth factor systems, and other biological systems with the same structure, behave similarly.

Growth factors are well-established proto-oncogenes, and understanding the conditions under which their production is evolutionarily stable in tumors is directly relevant to therapeutic strategies aimed at disrupting intra-tumor cooperation (Lemmon & Schlessinger, 2010; Greaves & Maley, 2012; Archetti & Pienta, 2019). More broadly, the metabolic cost of secreted protein production is rarely measured directly, yet it determines whether production is evolutionarily stable in mixed populations — a consideration relevant to both cancer biology and the design of engineered cell lines for recombinant protein production.

## Methods

### Generation of producer and non-producer cell lines

Two lentiviral constructs were used to generate the cell lines used in this study: pLVX-TetOne-rmIL3-HygR (TetOne-IL3; cloned by Genscript and sequence-verified), which drives doxycycline-inducible expression of murine IL3, and pLVX-PGK-Puro-IRES-GFP (PIG; Addgene), which constitutively expresses GFP and a puromycin resistance cassette. Lentiviral particles were produced by transfecting HEK293T cells (ATCC) with each pLVX construct together with third-generation lentiviral packaging plasmids and VSV-G envelope plasmid by calcium phosphate transfection. Viral supernatant was collected at 24 h and 48 h post-transfection, pooled, and concentrated by polyethylene glycol (PEG) precipitation. To generate producer cells, Ba/F3 cells were transduced with TetOne-IL3 and stably expressing clones selected with hygromycin. To generate fluorescently labeled non-producer cells for co-culture tracking experiments, Ba/F3 cells were transduced with PIG and selected with puromycin.

### Cell culture and growth tracking

Ba/F3 cells (DSMZ) were maintained in RPMI-1640 (Sigma-Aldrich) supplemented with 10% fetal bovine serum (Corning) and 1% penicillin–streptomycin (Life Technologies) at 37°C in a humidified incubator with 5% CO_2_. Routine maintenance medium was supplemented with 1 ng ml^−1^ recombinant murine IL3 (R&D Systems, #403-ML); producer cells were maintained without exogenous IL3 unless otherwise stated.

For pure-population growth rate measurements, 50,000 producer or non-producer cells were seeded per well in 6-well plates in triplicate in medium with or without exogenous IL3. For co-culture competition experiments, producer and non-producer cells were mixed at ratios spanning 0% to 100% producers and seeded at 500,000 cells per well in medium containing 0 to 1 ng ml^−1^ IL3, as specified for each experiment. Cell counts, viabilities, and GFP^+^ non-producer fractions were measured daily by flow cytometry (BD Accuri C6). Briefly, approximately 100 µl of cell suspension was removed from each well, transferred to a 1.5 ml microcentrifuge tube, and analyzed. The remaining suspension was centrifuged at 1,000 × *g* for 5 min, the supernatant aspirated, and the cell pellet resuspended in fresh medium at the appropriate IL3 concentration. For subculturing, a defined fraction of the resuspended cells was transferred to a new plate; this fraction was recorded and used to correct subsequent cell count measurements for dilution.

### Mathematical model

The fitness W_*i*_ of producers (*i*=C, cooperators) and non-producers (*i*=D, defectors) are given by

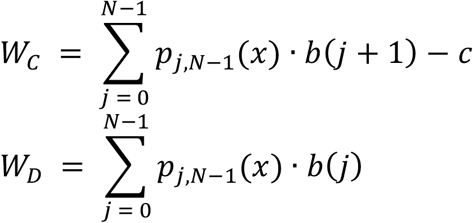

Respectively, where *x* is the frequency of producers in the population and

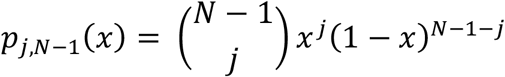

is the probability mass function of a binomially distributed random variable *j* with parameters (*N*−1, *x*) and the benefit function

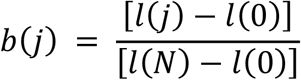

is the normalized version of the logistic function

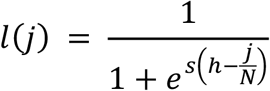

where the parameter *h* defines the position of the inflection point (*h*→1 gives increasing returns and *h*→0 diminishing returns) and *s* defines the steepness of the function at the inflection point (*s*→∞ models a threshold public goods game; *s*→0 models linear (additive) benefit; the normalization prevents the logistic function from becoming constant for *s*→0). If the benefit function is linear, these can be simplified to:

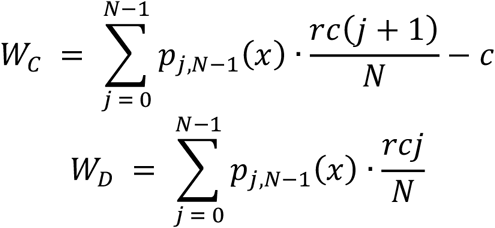

The replicator dynamics is

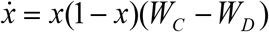

Where the gradient of selection is

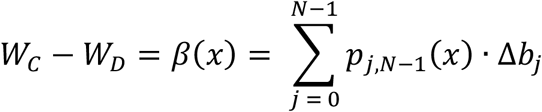

a polynomial in Bernstein form (Archetti 2018), with Bernstein coefficient

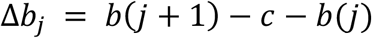

Beyond the two trivial rest points *x* = 0 and *x* = 1, further interior rest points are given by *β*(*x*)=0. The fitness of mixed population is given by *x*·W_C_+(1-*x*)·W_D_.

